# A dynamic pattern of local auxin sources is required for root regeneration

**DOI:** 10.1101/783480

**Authors:** Rotem Matosevich, Itay Cohen, Naama Gil-Yarom, Abelardo Modrego, Carla Verna, Enrico Scarpella, Idan Efroni

**Author notes:** Corresponding author: Idan Efroni.

## Abstract

Following removal of its stem cell niche, the root meristem can regenerate by recruitment of remnant cells from the stump. Regeneration is initiated by rapid accumulation of auxin near the injury site but the source of this auxin is unknown. Here, we show that auxin accumulation arises from the activity of multiple auxin biosynthetic sources that are newly specified near the cut site and that their continuous activity is required for the regeneration process. Auxin synthesis is highly localized and PIN-mediate transport is dispensable for auxin accumulation and tip regeneration. Roots lacking the activity of the regeneration competence factor *ERF115*, or that are dissected at a zone of low-regeneration potential, fail to activate local auxin sources. Remarkably, restoring auxin supply is sufficient to confer regeneration capacity to these recalcitrant tissues. We suggest that regeneration competence relies on the ability to specify new local auxin sources in a precise spatio-temporal pattern.

## Introduction

Continuous plant growth is supported by the activity of meristems, a complex, multi-tissue organ that houses the stem cell niche (SCN), which in turn is organized around a group of cells with low mitotic activity called the quiescent center (QC) (Heidstra and Sabatini, 2014)). Remarkably, ablation or excision of the SCN, including the QC, results in complete meristem regeneration within a few days (Feldman, 1976; Reinhardt et al., 2003; Xu et al., 2006; Sena et al., 2009; Efroni, 2018). This process is best studied in the root meristem, where excision of the entire root tip triggers rapid *de novo* specification of a SCN from multiple remaining differentiated cells at the stump (Efroni et al., 2016). The process is controlled by regeneration competence factors, such as *ERF115* (Heyman et al., 2016; Zhou et al., 2019), as well as multiple plant hormones (Efroni et al., 2016; Heyman et al., 2016; Zhou et al., 2019).

Key to root tip regeneration is the activity of auxin (Sena et al., 2009; Efroni et al., 2016). (indole-3-acetic acid; IAA), one of the main phytohormones in plants which plays a role in many plant developmental processes (Weijers and Wagner, 2016). Auxin is synthesized from tryptophan via *TRYPTOPHAN AMINOTRANSFERAS OF ARABISOPSIS 1 (TAA1)*, producing indole-3-pyruvate, which then undergoes decarboxylation by *YUCCA (YUC)*(Zhao, 2010). In its reduced form, IAA can enter the cell either via diffusion or via the influx carriers *AtAUX1-LAX*. However, its efflux is mainly mediated by *PIN-FORMED (PIN)* family transporters which are polarly distributed on the plasma membrane and regulate auxin distribution in plant tissues (Adamowski and Friml, 2015). In the root, auxin is produced in the QC (Petersson et al., 2009; Brumos et al., 2018) and is transported via the PINs to generates a concentration maxima at the SCN (Grieneisen et al., 2007). This maxima serves as an instructive signal for the specification of the distal part of the root (Sabatini et al., 1999). During embryogenesis, PINs transport auxin from multiple marital or embryo-internal sources to its development-relevant position (Robert et al., 2013; Robert et al., 2015).

How auxin concentration is regulated during root regeneration and what is its function in the process is unclear. Following injury, PIN polarization in the stump remained unaltered and while application of the auxin efflux inhibitor 1-N-naphthylphthalamic acid (NPA) (Petrásek et al., 2003) disrupted root meristem regeneration, some growth at the injured tip was still apparent (Sena et al., 2009). Additionally, while injury to plant tissue is often associated with a broad increase in auxin biosynthetic activity (Sztein et al., 2002; Chen et al., 2016; Druege et al., 2016; Xu et al., 2017), the contribution of auxin biosynthesis to auxin accumulation and distribution is unknown.

Here, we utilize tissue and stage-specific inhibition of the auxin biosynthetic machinery in order to dissect the dynamics role of local auxin biosynthesis during tip regeneration. Based on this characterization, we analyze recalcitrant tissue and show that regeneration failure stems from the inability to activate local auxin sources at appropriate temporal sequence.

## Results

### Auxin biosynthesis is required for root tip regeneration

To test the role of auxin biosynthesis in root tip regeneration, we first examined mutants of YUC3/5/7/8/9, the main YUC enzymes acting in the root. Quintuple *yuc3 yuc5 yuc7 yuc8 yuc9* (*yucQ)* mutants have short roots with very small meristems (Chen et al., 2014a). Regeneration rates are affected by meristem size, so in order to measure regeneration rates in this background, *yucQ* were grown on 5 nM IAA-supplemented plates, which rescued the growth and meristem defects (Chen et al., 2014a). Tips were then cut and seedlings transferred to an IAA-free medium, along with WT controls. Three days post-cut, roots were scored for regeneration of a new root tip. Regeneration rates in *yucQ* mutants were reduced compared to WT, an effect that could be recapitulated by post-cut treatment of regenerating roots with YUCCASIN-DF, a competitive inhibitor of YUC (Tsugafune et al., 2017) (Fig. 1A). Similar treatment with the TAA inhibitor L-Kyn (He et al., 2011), led to a dose-dependent inhibitory effect on regeneration, with the highest concentration leading to complete abortion of regeneration, indicating that auxin biosynthesis is required for root tip regeneration (Fig. 1A). Due to its strong phenotype and complete loss of regeneration capacity, we chose to utilize L-Kyn treatment to further probe the role of auxin biosynthesis in regeneration.

**Figure 1.**
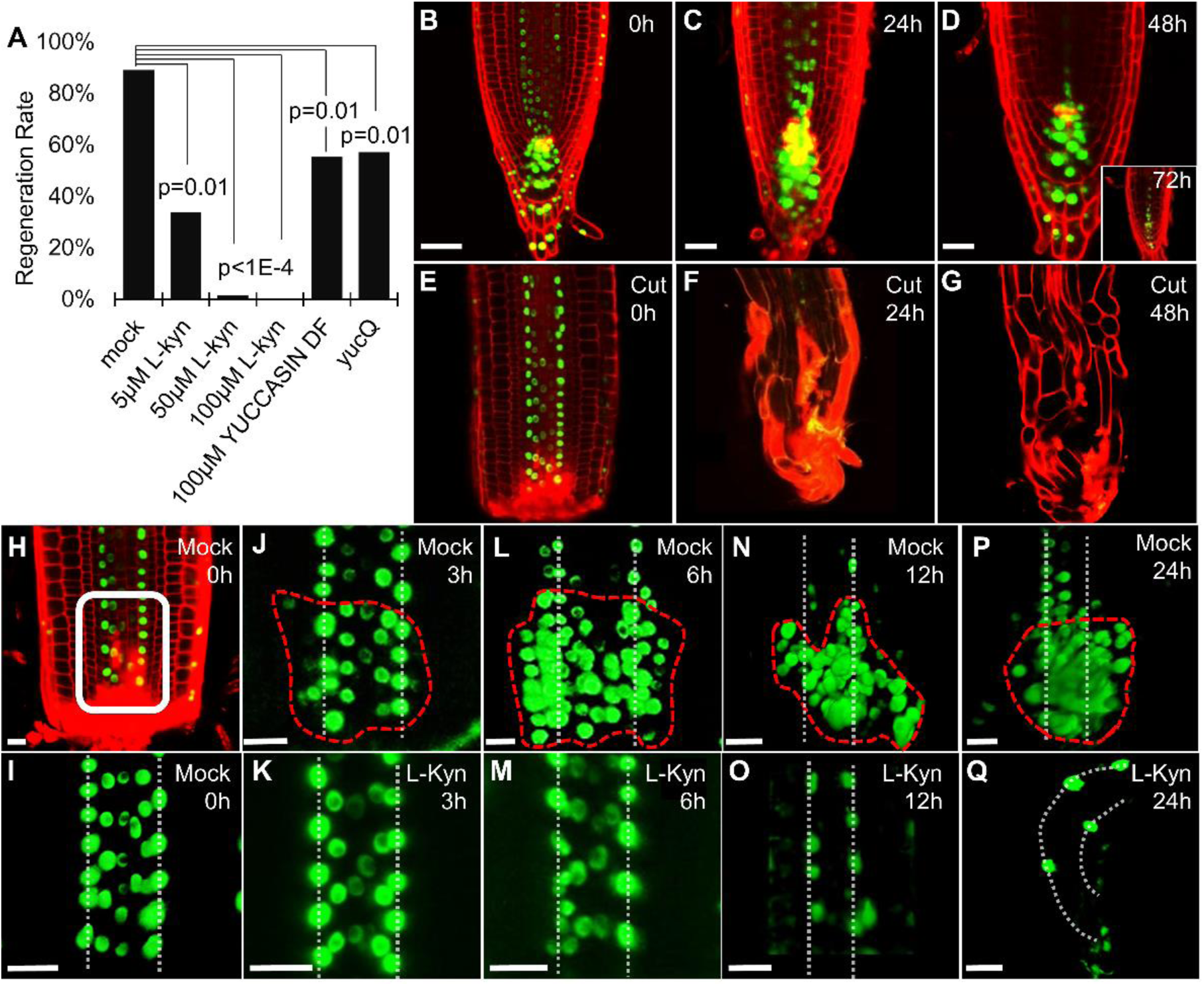
Auxin biosynthesis is required for root tip regeneration. **A)** Root tip regeneration rates in plants with disrupted auxin biosynthesis activity, and either treated with different concentrations of L-Kyn or 100µM YUCASIN DF or mutant for 5 YUC genes (*yucQ*) (shown p-values are for Tukey HSD on a logistic regression model; n=55, 62, 65, 75, 47, 63 for mock, 5µM L-Kyn, 50µM L-Kyn, 100µM L-Kyn, 100µM YUCASIN DF and yucQ, respectively). **B-G)** Confocal time-series of 100µM L-Kyn-treated *DR5rev:3xVENUS-N7* plants showing uncut (B-E) or regenerating roots (E-G). **H-Q)** Expression of *pDR5rev:3XVENUS-N7* immediately after dissection (H-I) or during regeneration (J-Q) of tips treated with mock (J,L,N,P) or 100µM L-Kyn (K,M,O,Q). White box in (H) marks the area shown in (I-Q). Dotted vertical lines mark the protoxylem. Dashed red lines mark the forming auxin peak. Propidium iodide was used to stain cell walls (red). Scale bars are 50µm (B-G) and 20µm (H-Q).

In uncut roots, L-Kyn treatment caused meristems to gradually shrink over a period of several days (Brumos et al., 2018), but after 3 days of L-Kyn treatment, meristematic cells were still present at the root tip and the expression of the auxin response marker DR5 retained its normal expression pattern, albeit with weakened intensity (Fig. 1B-D). In strike contrast, cut root tip were extremely sensitive to L-Kyn treatment and after just 24h of treatment, no meristem was apparent and cells appeared fully differentiated (Fig. 1E-G). High-resolution monitoring of *DR5*, one of the earliest markers of regeneration (Ulmasov et al., 1995; Sena et al., 2009; Efroni et al., 2016), showed remnant DR5 expression in the xylem immediately after the cut (Fig. 1H-I). By 3 hours post-cut (hpc), cells immediately adjacent to the protoxylem began to express DR5 (Fig. 1J) but no such induction was apparent in L-Kyn-treated roots (Fig. 1K). By 6 hpc, the DR5 signal expanded laterally from the protoxylem and gradually became restricted to the basal part of the stump, forming a distinct auxin peak by 12-24 hpc. L-Kyn treatment completely inhibited this expansion and DR5 expression remained limited to the xylem, until the root meristem differentiated at ∼24 hpc (Fig. 1I-Q; Fig 1E). The lack of early *DR5* activation in L-Kyn treated cut roots suggests that auxin biosynthesis normally occurs shortly following tip dissection and is responsible for establishment of a new auxin maxima near the cut site.

### Continuous auxin biosynthesis is required to prevent premature differentiation and for pattern recovery during regeneration

To determine whether auxin synthesis is required just to “kick-start” regeneration or whether it is required throughout the process, roots were dissected and allowed to recover on mock plates for varying times prior to or post L-Kyn treatment. An immediate but short (6h) treatment with L-Kyn was associated with a dramatic reduction in regeneration rate, while roots treated for 24h failed to recover altogether. Remarkably, treatment with L-Kyn at different time points during regeneration, up until ∼72 hpc, resulted in regeneration failure, an effect that was mitigated upon addition of IAA to the L-Kyn treatment medium (Fig. 2A). This confirms that the inhibition of auxin biosynthesis, rather than other possible non-specific side effects of L-Kyn, is critical for regeneration and indicates that auxin biosynthesis is required both for early and late stages of regeneration.

**Figure 2.**
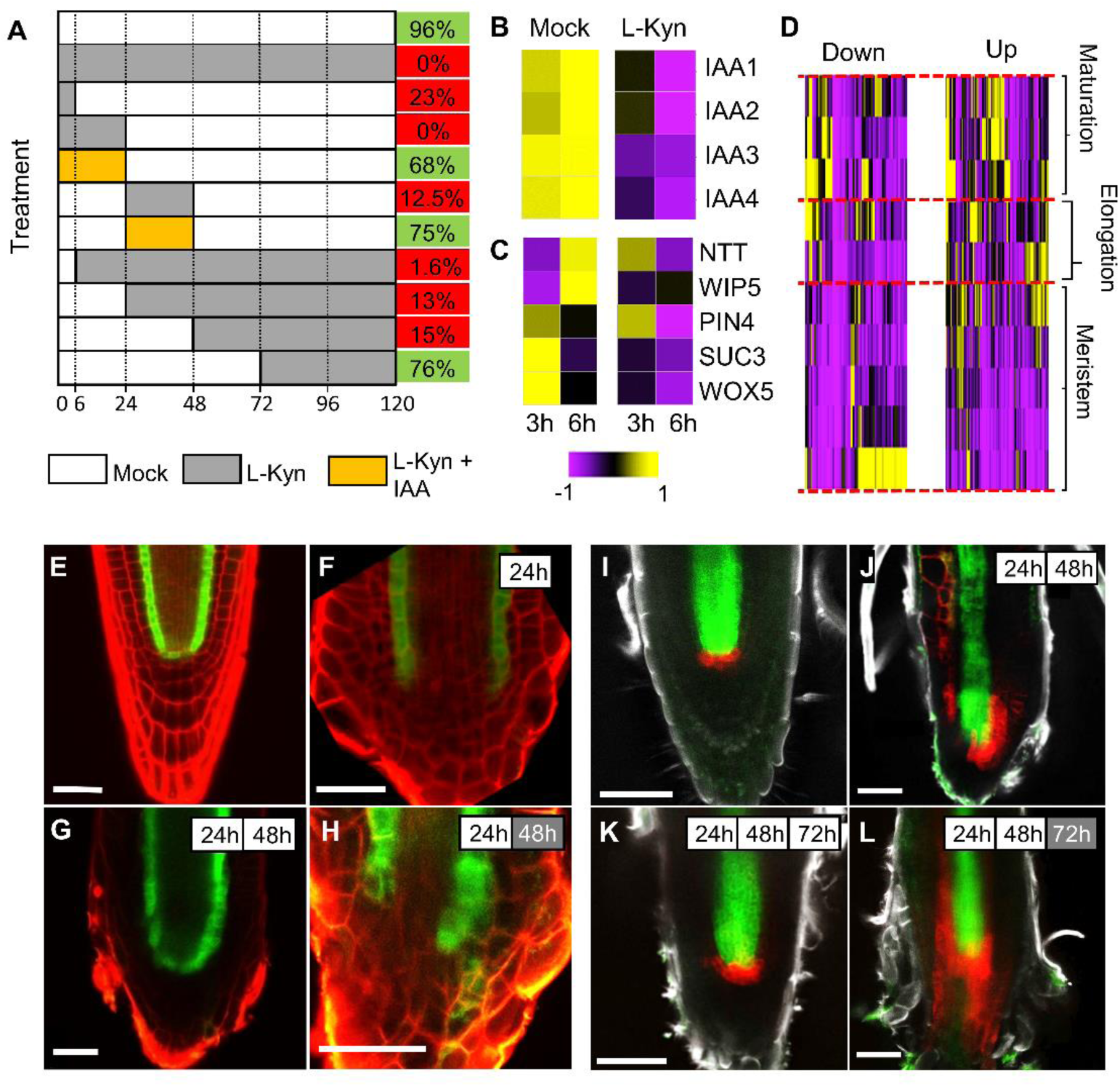
Auxin biosynthesis is required throughout the regeneration process. **A)** Regeneration rates of 7 DAS roots at different time points of mock, 100µM L-Kyn or 100µM L-Kyn+100nM IAA treatment. Regeneration rates are shown on the right (Green marks rates of 66% and up, red is lower than 66%). **B-C)** Expression of auxin response genes (B) and regeneration-induced genes (C). **D)** Expression of genes modified by 6h L-Kyn treatment at different regions of the root. **E-L)** Confocal images of SCR:YFP (E-H) or WOX5:mCherry x WOL:GFP (I-L) before cut (E,I) or during regeneration, upon mock (F-G, J-K) or 100µM L-Kyn (H,L) treatment, at the specified time points. White and gray boxes mark time spent on mock or L-Kyn plates, respectively. Scale bars: 50µm.

In order to understand the specific role of early auxin synthesis a transcriptomic time-series of L-Kyn-treated root tip regeneration was generated by immediately transferring dissected roots to L-Kyn or mock agar plates for 3 h or 6 h, after which the root meristem stump was isolated and profiled. As a control, intact meristems treated with L-Kyn for 3h or 6h were isolated and profiled (Supplemental Fig. 1A; Supplemental Table 1). Consistent with the requirement for auxin biosynthesis in early stages of post-cut regeneration, the increased expression of the auxin response genes IAA1/2/3/4 (Weijers and Wagner, 2016) was largely blocked by L-Kyn treatment (Fig. 2B). Further, induction of previously identified early markers of regeneration (Efroni et al., 2016), was attenuated at 3 h of L-Kyn treatment and fully lost at 6 h, indicating early termination of the regeneration process (Fig. 2C). To identify the processes affected by auxin biosynthesis during regeneration, we first selected only the genes that were specifically modified by L-Kyn in the regenerating roots, as compared to the uncut meristem (Supplemental Fig. 1B; Supplemental Table 1). GO analysis revealed that genes suppressed by the 3 h L-Kyn treatment were enriched for wound response, histogenesis and auxin response, while genes suppressed following 6 h exposure were highly enriched for cell cycle and cytokinesis processes, suggesting that auxin synthesis is required to induce early cell division in the stump (Supplemental Fig. 1C-D). Mapping of the genes modified by 6 h L-Kyn treatment onto the gene expression map of the root (Brady et al., 2007) revealed that genes downregulated by L-Kyn treatment were generally enriched in the meristematic zone, especially in the stem-cell-niche region, consistent with L-Kyn inhibition of the formation of a new SCN. In contrast, genes induced by L-Kyn were normally expressed at higher parts of the meristem and in the elongation zone (Fig. 2D). Taken together, these changes indicate that without auxin biosynthesis, cells in the stump are unable to activate cell cycle programs and seem to undergo precocious differentiation.

Later stages (12-72hpc) of regeneration are marked by gradual repatterning of the regenerating tip (Sena et al., 2009; Efroni et al., 2016). The endodermis/QC marker *SCARECROW:YFP (SCR:YFP)* was monitored to determine whether auxin synthesis is required just for cellular proliferation or also for tissue pattern recovery. During normal regeneration, expression of tissue identity markers is lost near the cut site at 0-24 hpc, and then gradually regained (Efroni et al., 2016). When *SCR:YFP* plants were allowed to recover for 24 h and then treated with L-Kyn for an additional 24 h, roots did not establish normal endodermis patterning and resembled roots at 24 hpc stage (Fig. 2E-H), suggesting that at 24-48 hpc, auxin biosynthesis is required for proper cell identity recovery. To determine whether the biosynthesis is also required for patterning during later stages of regeneration, we examined the recovery of *WOX5*, which is normally confined to the QC cells, but during regeneration is broadly expressed in the vicinity of the stele until ∼72 hpc, when it is regains its proper localization (Fig. 2I-K). Consistent with the role of biosynthesis in repatterning of injured tissue, when *WOX5:mCherry* roots were transferred to L-Kyn following 48 h of recovery, localized QC expression was not established and its expression remained diffused (Fig. 2L). In conclusion, auxin biosynthesis is required throughout the regeneration process both for activation of cell proliferation and for pattern recovery, serving different roles at different stages.

### Multiple local auxin sources are sequentially induced during regeneration

Auxin is classically thought to be synthesized at the shoot and transported to the root. However, recent evidence suggests that auxin is synthesized much closer to its region of accumulation (Chen et al., 2014a; Brumos et al., 2018). To identify whether the source of auxin required for regeneration is local or remote, cut roots were placed on split plates containing L-Kyn in both sides of the plate and IAA supplied either to the top part of the plant (shoot and top part of the root), or directly to the regenerating tip. Regeneration was restored only when the tip was located in the IAA-containing part of the plate (Fig. 3A), indicating that auxin is likely produced locally during regeneration.

**Figure 3.**
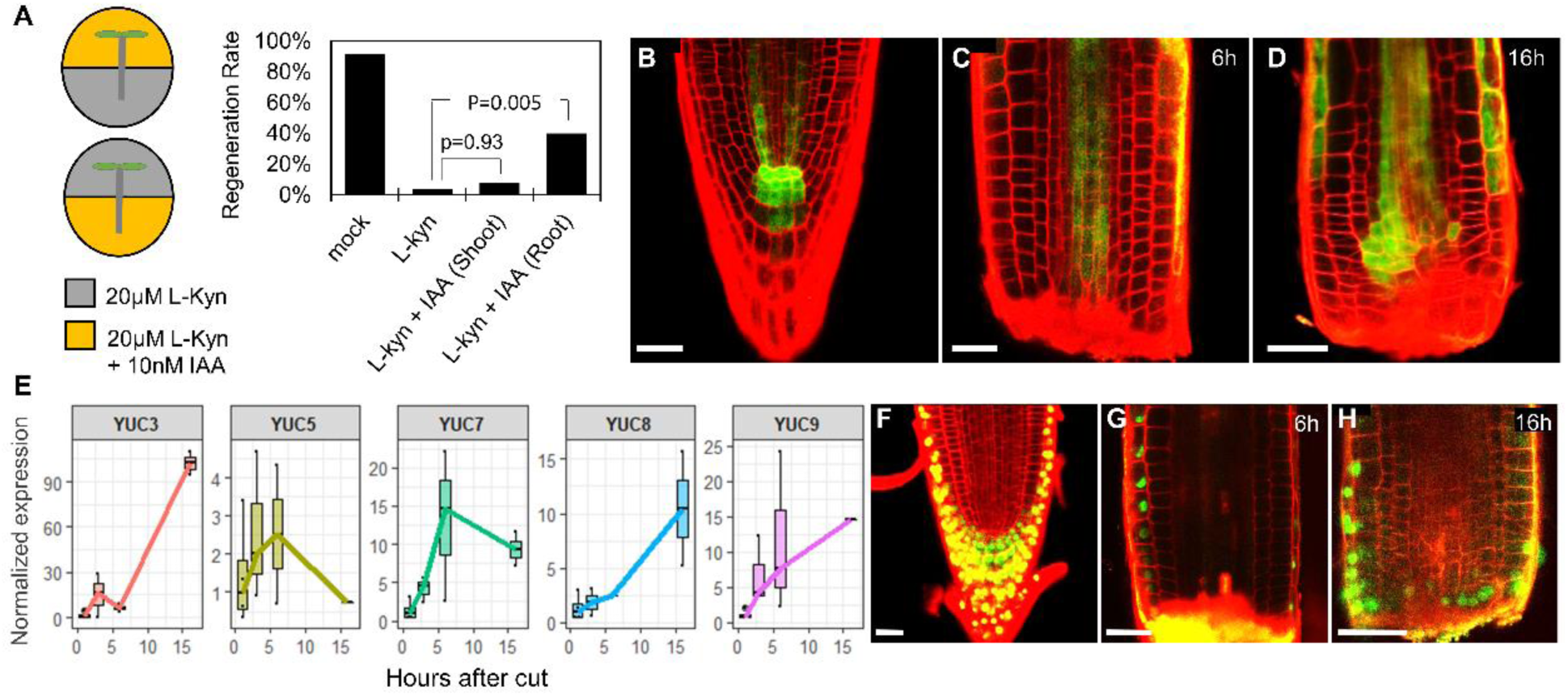
Local auxin synthesis by multiple sources during root tip regeneration. **A)** Regeneration rates of 7 DAS roots treated with 20µM L-Kyn and supplemented with 10nM IAA either at the shoot and upper part of the root or at the lower part of the root (shown p-values are for Tukey HSD on a logistic regression model; n=50, 50, 47, 50 for mock, 20µM L-Kyn, 20µM L-Kyn+10nM IAA shoot, 20µM L-Kyn+10nM IAA root, respectively). **B-D)** Confocal images of uncut (B) or regenerating roots (C-D) expressing *TAA1p:GFP-TAA1*. **E)** qRT-PCR measurement of YUC genes in isolated meristems of regenerating roots at different time points. Expression was normalized to that of cut meristems isolated immediately after cut. **F-H)** Confocal images of uncut (F) or regenerating roots (G-H) expressing *pYUC9:VENUS-termYUC9*. Scale bars: 20µm.

During steady-state root meristem growth, auxin is mainly synthesized in the QC (Brumos et al., 2018), which is fully removed upon root tip dissection. To determine the source of auxin during regeneration, we examined the temporal expression of its biosynthetic enzymes TAA and YUCCA. *TAA1:TAA1:GFP* was found to be expressed in the protoxylem at early stages of regeneration (Fig. 3B-D), consistent with biosynthesis being required for the early appearance of the DR5 signal next to the xylem (Fig. 1J-K). qRT-PCR of isolated regenerating meristems at 0hpc, 3hpc, 6hpc and 16hpc, detected rapid induction of five YUC genes in the regenerating root (Fig. 3E). Curiously, the *YUC* exhibited different expression dynamics, with YUC5/7 exhibited transient expression, while the expression of YUC3/8/9 gradually increased until 16 hpc. Analysis of the expression of a reporter for YUC9, (*pYUC9:VENUS-termYUC9)*, which is normally confined to the root cap (Fig. 3F), revealed that its expression at 16 hpc was localized to the immediate vicinity of the cut site (Fig. 3G-H). This localized induction was consistent with the enhanced auxin accumulation above the cut site between 12-24 hpc (Fig. 1N,P; (Efroni et al., 2016)).

The gradual establishment of DR5 expression, first at the vicinity of the protoxylem and then at the cut site region, together with the induction of multiple YUC enzymes, suggests activation of multiple distinct auxin sources during regeneration. To test this hypothesis, we generated an artificial miRNA targeting YUC2/3/5/6/7/8/9 (*amirYUC)* and expressed it using different promoters active during regeneration. Expression of *amirYUC* under the QC-specific WOX5 promoter, exhibited disorganized QC cell divisions in 9 DAS plants (Fig. 4A-B), but despite *WOX5* broad expression in ground tissue during regeneration (Fig. 2J-K; (Efroni et al., 2016)), *WOX5:amirYUC* plants did not exhibit a significant reduction in regeneration, suggesting that auxin is not provided to the regenerating meristem by a re-established QC (Fig. 4E). To test whether the xylem serves as an auxin source during regeneration, we expressed *amirYUC* under the *AHP6* promoter, which is expressed in the protoxylem and xylem-pole-pericycle before and immediately after the cut, but gradually loses its expression near the cut site by ∼24 hpc (Efroni et al., 2016). *AHP6:amirYUC* plants exhibited mild disruption to QC cell division patterns (Fig. 4C) and in agreement with the xylem serving as an auxin source during regeneration, caused a significant reduction in regeneration rate (Fig. 4E). Finally, to test whether an auxin source near the cut site plays a role in regeneration, *amirYUC* was expressed under the pYUC9 promoter, which in intact meristems, is excluded from the QC (Fig. 3F). *YUC9:amirYUC* resulted in mild disruption to the cellular organization of the distal meristem (Fig. 4D) and to rare pin-like inflorescence meristems (Supplemental Fig. 2A-B), a phenotype previously observed for *yuc1 yuc4 npy1* (Cheng et al., 2007). Remarkably, *YUC9:amirYUC* had low regeneration rates, comparable to the *yucQ* mutants (Fig. 5E; Fig. 1A), indicating that this auxin source, possibly acting near the cut site, plays a significant role in regeneration and that, overall, local and dynamic auxin biosynthesis is required to initiate and sustain root regeneration.

**Figure 4.**
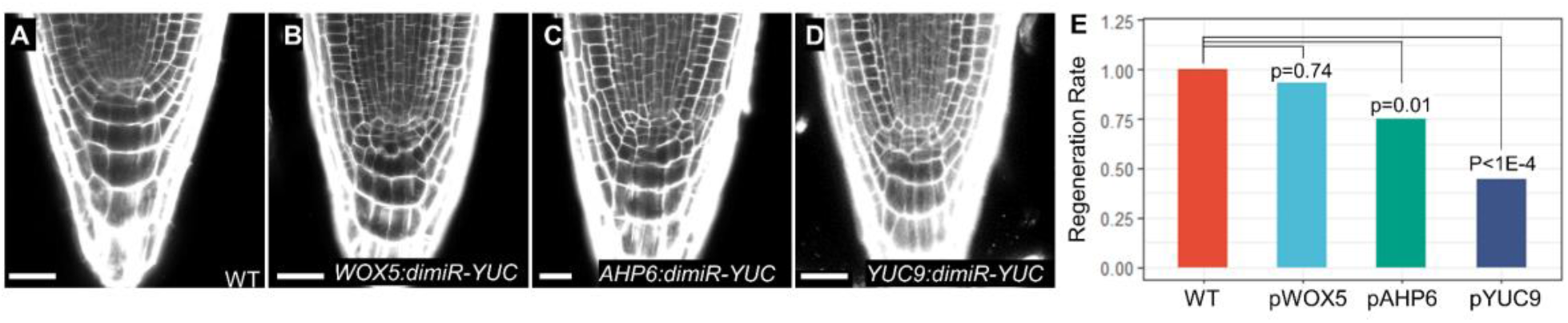
Stage- and tissue-specific knockdown of YUC during regeneration. **A-D)** Confocal images of uncut 9 DAS meristem of WT plants (A) or mutants expressing *amiRYUC* under different tissue-specific promoters (B-D). **E)** Regeneration rates of mutants expressing *amiRYUC* under different tissue-specific promoters (shown p-values are for Tukey HSD on a logistic regression model; n=154, 60, 68, 23 for WT, *WOX5:amiRYUC, AHP6:amiRYUC*, and *pYUC9:amiRYUC* respectively). Scale bar: 50µm.

**Figure 5.**
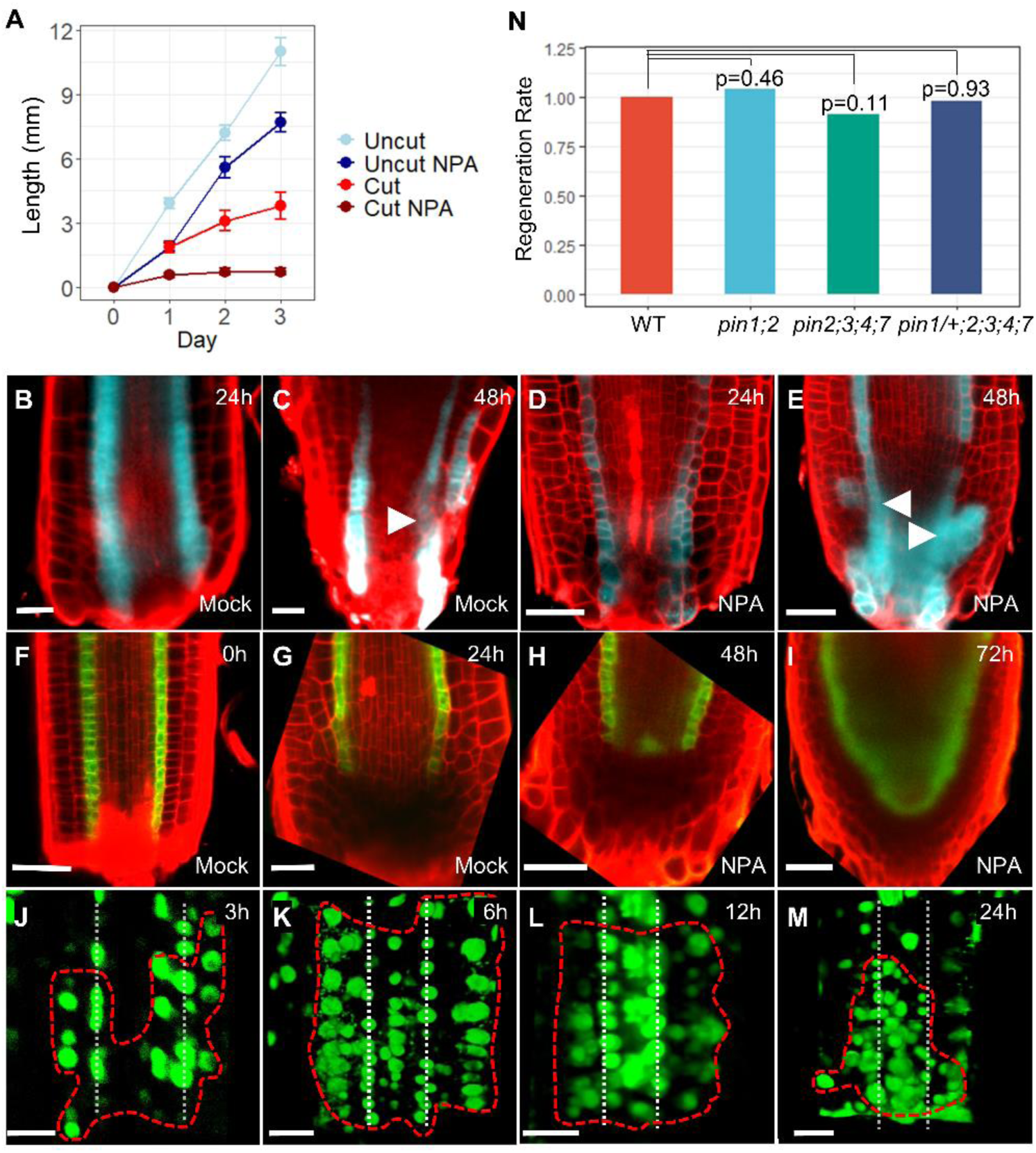
PIN-mediated polar auxin transport is not required for regeneration. **A)** Growth of uncut and cut roots transferred at 7 DAS to control or 10µM NPA-supplemented plates. **B-I)** Confocal images of regenerating root tips carrying lineage CFP-marked clones induced by the *SCR* promoter (B-E) or expressing the labile *SCR:YFP* reporter (F-I), allowed to regenerate on mock (B-D, F-G) or 10µM NPA-supplemented plates (D-E, H-I). Arrowheads mark the origin of lateral divisions. **J-M)** Close-up of the regeneration region of *DR5rev:3xVENUS-N7* plants treated with 10µM NPA. Dotted vertical lines mark the protoxylem; compare to Fig. 1H-Q. **N)** Regeneration rates of *pin* mutants. No significant change in regeneration rate was observed (p-values are for Tukey HSD on a logistic regression model; n=74,74,85, 28 for WT, *pin1;2* and *pin2;3;4;7*, and *pin1/+;2;3;4;7*, respectively). Scale bar: 50µm.

### PIN-mediated polar auxin transport is not required for root tip regeneration

During steady state root growth, PINs mediated transport act together with local biosynthesis to maintain the root meristem (Brumos et al., 2018). Similarly, during root regeneration from leaves, auxin synthesis is induced in the leaf, which the PINs are responsible to direct the auxin to the site of root initiation (Chen et al., 2016). Blocking polar auxin transport using NPA was reported to arrest root tip regeneration (Sena et al., 2009) and to better understand the possible effect of PINs on root tip regeneration, we revisited these experiments. As previously reported, NPA treatment caused complete growth arrest of regenerating roots (Fig. 5A). However, close examination revealed that proliferation at the cut root tip was not inhibited and that by 72 hpc, the regenerating roots resembled uncut NPA-treated roots (Supplemental Fig. 3A-D), suggesting that regeneration may occur even with NPA treatment.

To verify that the growth of the tip under NPA represents true regeneration, we used a lineage tracing system to identify whether a new SCN is formed. The new SCN is specified at 24-48 hpc and is marked by a transition from an endodermal cell to an epidermis stem cell, made evident by lateral cell divisions originating from the endodermis (Fig 5C-D; (Efroni et al., 2016)). NPA treatment did not disrupt the appearance of these lateral divisions and they originated from similar positions as those occurring in mock-treated roots (Fig. 5C-F). Further consistent with proper regeneration, expression pattern recovery of the QC/endodermal marker *SCR:YFP* and the ground tissue marker J0571 were not affected by NPA treatment, other than a mild delay (Fig. 5G-J; Supplemental Fig. 5E-L). In agreement, expression of DR5 was induced next to protoxylem cells at 3 hpc even when treated with NPA (Fig. 5K) and while by 6 hpc and 12 hpc, NPA-treated roots exhibited a broader DR5 expression domain than control plants, by 24 hpc, the peak was confined to the distal end of the root tip, similar to control (Fig. 5L-M; Fig. 1N,P).

To genetically verify the PINs are not required for root tip regeneration, we tested the regeneration capacity of plants with mutant PIN transporters PIN1/2/3/4/7, which perform partially redundant functions in the root (Blilou et al., 2005). Regeneration rates and meristem morphology of *pin1 pin2, pin2 pin3 pin4 pin7* and *pin1/+;2;3;4;7* were not significantly different than wild type plants (Fig. 5N; Supplemental Fig. 4). Quintuple *pin1;2;3;4;7* plants germinated normally, but produced short roots with very small meristems (Supplemental Fig. 4) which did not allow for measurement of regeneration rates. However, when carefully cut, regeneration was observed, and the newly formed tip resembled that of NPA-treated regenerating plants (compare Supplemental Fig. 4K with Supplemental Fig. 1D). Finally, assessment of regeneration in mutants in the callosin-like protein *BIG*, which are defective in auxin transport (Gil et al., 2001), showed a new auxin peak formed near the cut site at 6 hpc, similar to regeneration in WT plants (Supplemental Fig. 5). Taken together, genetic and pharmacological evidence suggest that PIN transporters may act to refine auxin distribution during regeneration, but are required neither for auxin peak formation, nor for its distal localization nor for regeneration *per se*.

### The regeneration competence factor *ERF115* is required for activation of the early auxin response

Given that active local auxin biosynthesis is required for root tip regeneration, we hypothesized that factors that regulate regeneration competence may involve, directly or indirectly, in the activation of the auxin biosynthetic machinery. One of the key factors in determining regeneration competence is the AP2-like protein *ERF115*, which is rapidly induced in the vicinity of the cut site (Fig. 6A) and expression of its dominant negative *35S:ERF115-SRDX* caused a severe reduction in regeneration capacity (Heyman et al., 2016; Zhou et al., 2019). While *ERF115* was induced near the cut site even when auxin biosynthesis was inhibited using L-Kyn (Fig. 6B), *ERF115-SRDX* plants which fail to regenerate, did not generate a normal auxin signaling peak at 6 hpc (Fig. 6C). Consistently, induction of auxin biosynthesis enzymes was severely attenuated in this mutants, and only YUC3 and YUC7 were induced by 6 hpc, with their expression rapidly lost by 16 hpc (Fig. 6D). Remarkably, application of 5nM IAA to the root tip immediately after dissection of *35S:ERF115-SRDX* plants, resulted in almost complete recovery of root regeneration potential (Fig. 6E-F), suggesting that the inability to activate biosynthesis in these mutants is responsible for the loss of regeneration capacity.

**Figure 6.**
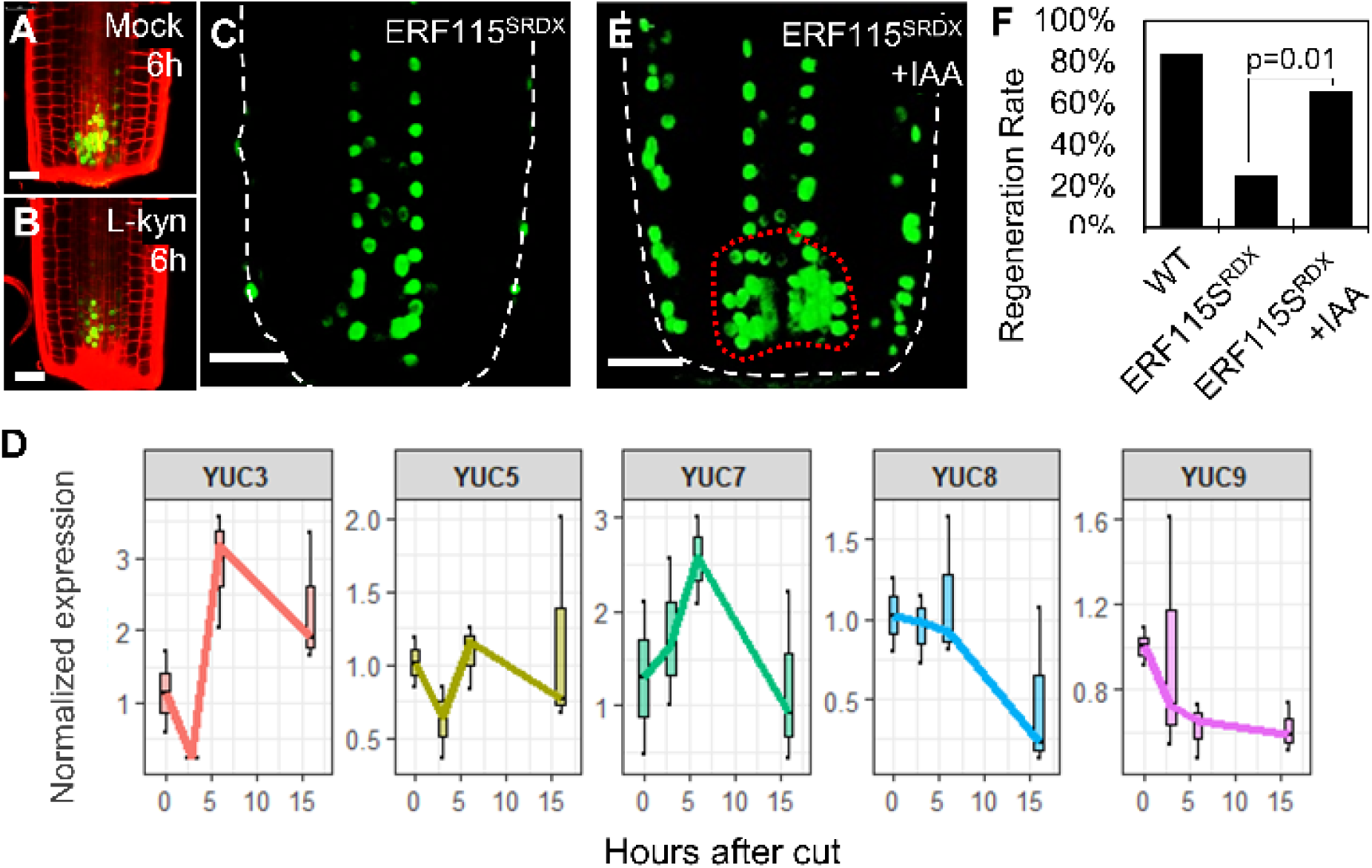
Auxin synthesis in regenerating *ERF115* dominant negative plants. **A-B)** Confocal images of *ERF115:GFP* in regenerating roots treated with mock (A) or 100µM L-Kyn (B). **C)** Confocal images of *35S:ERF115-SRDX DR5rev:3xVENUS-NLS* roots at 6 hpc. **D)** qRT-PCR measurement of YUC expression in isolated meristems of regenerating *35S:ERF115-SRDX* roots. Expression was normalized to that of isolated meristems immediately after cut. **E)** Confocal images of *35S:ERF115-SRDX DR5rev:3xVENUS-NLS* root treated with 5nM IAA at 6 hpc. **F)** Regeneration rates of *35S:ERF115-SRDX* 7 DAS roots on mock or 5nM IAA (p-values are for Tukey HSD on a logistic regression model; n=115, 181, 182 for WT, *35S:ERF115-SRDX* on mock and *35S:ERF115-SRDX* on 5nM IAA, respectively). Scale bar: 50µm.

### Regulation of auxin levels is required and sufficient to confer regeneration competence to differentiated cells

Wild type plants exhibit a gradient of regeneration competence within the meristem, with regeneration rates dropping for cuts performed high in the meristem (Sena et al., 2009; Zhou et al., 2019). To test whether the loss of capacity to activate auxin synthesis underlies this drop in regeneration competence, we cut roots at a distance of ∼220µM from the QC, at a zone where almost no regeneration occurs (Sena et al., 2009). Surprisingly, by 6 hpc, a normal auxin peak was apparent in high-cut roots (Fig. 7A). However, this expression was not sustained and by 12 hpc, DR5 expression was lost and subsequently, root tips differentiated (Fig. 7B). This suggests that while early sources of auxin may be active in high cut roots, late sources may not be induced. Indeed, qRT-PCR showed that *YUC5* and *YUC7* was rapidly upregulated but returned to base levels after several hours (Fig. 7C). Curiously, *YUC9* was strongly induced early on, but its expression returned to base levels, consistent with previous observations (Fig. 7C; (Xu et al., 2017)). This was in striking contrast to its temporal expression pattern in low-cut roots, which regenerate normally (Fig. 3E). Application of 5nM auxin to high-cut roots resulted in recovery of the 12 h auxin peak (Fig. 7D) and in a significant increase in regeneration rates (Fig. 7E), indicating that auxin is not only required but also sufficient to provide regeneration competence to high-cut roots.

**Figure 7.**
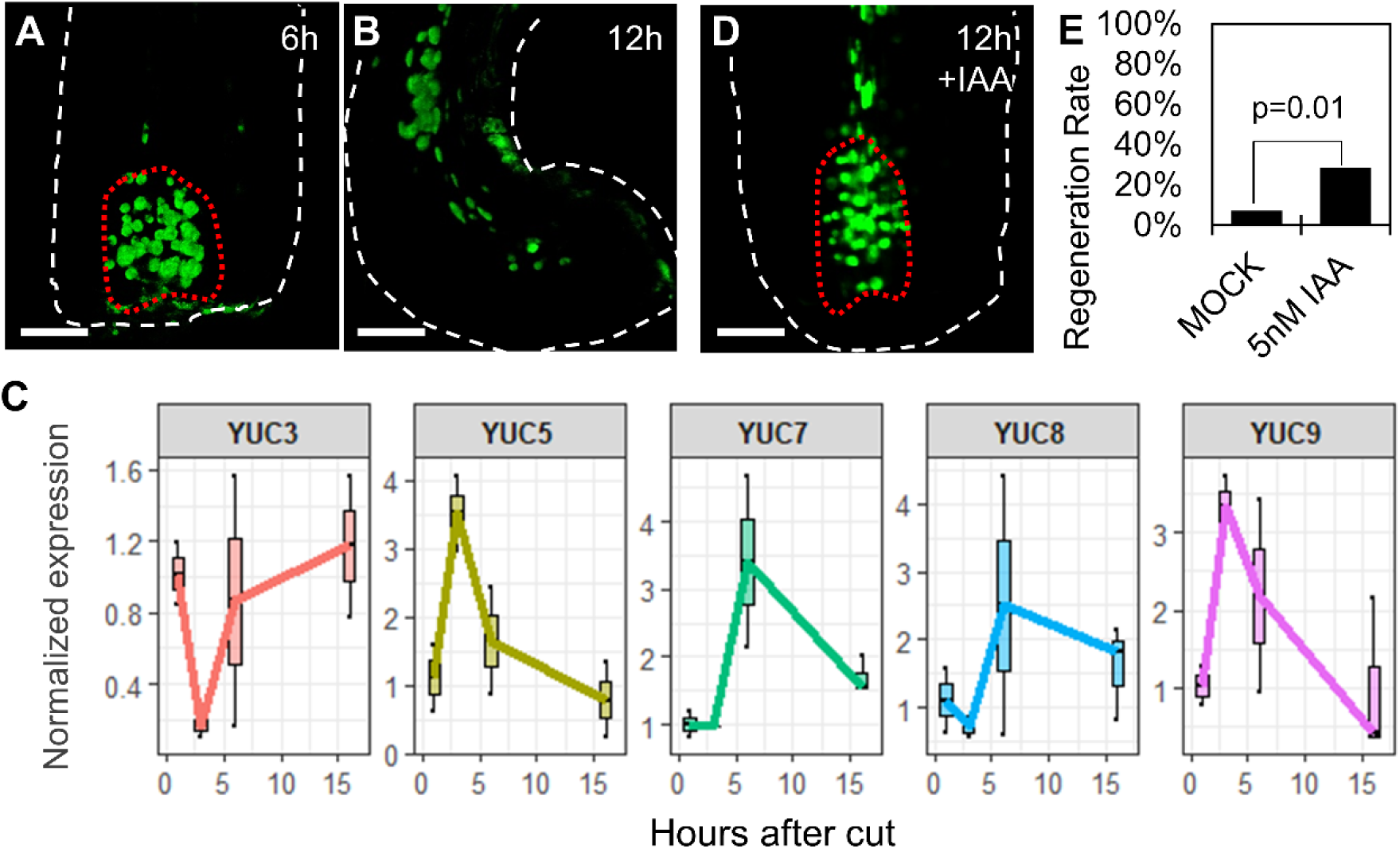
Auxin is sufficient to induce regeneration in high-cut roots. **A-B)** Confocal images of roots expressing *DR5rev:3xVENUS-N7* and cut at 220µm from the QC, at 6 hpc (A) or 12 hpc (B). **C)** qRT-PCR measurement of YUC expression in isolated meristems of regenerating roots cut at 220µm. Expression was normalized to levels in meristems immediately after cut. **D)** *DR5rev:3xVENUS-N7* of high-cut roots at 12 hpc treated with 5nM IAA. **E)** Regeneration rates of roots cut at 220µm from the QC and treated with 5nM IAA (χ-test; n=84, 84, for WT and 5nM IAA, respectively).

## Discussion

### Auxin synthesis is required and sufficient for regeneration competence

Regeneration competence in plants is widespread, but not universal, and it remains unclear why only some tissues are able to undergo complete regeneration. In the meristem, regeneration potential decreases with increasing heights of meristem injury (Sena et al., 2009; Zhou et al., 2019). Here, we show that the loss of competence in high-cut roots is due to the inability to specify and maintain new auxin sources. Interestingly, early auxin responses were induced in high-cut roots and were only lost at 12 hpc, suggesting that the cause for the inability to regenerate may not be in the immediate wound response, but rather in a long-term development program dictated by the internal state of cells. This capacity to activate temporal auxin synthesis programs may be a key universal feature of the “regeneration competence” of the tissue. Indeed, the ability of cut *Arabidopsis* leaves to produce microcalli structures and generate adventitious roots relies on auxin synthesis (Chen et al., 2014b; Chen et al., 2016; Bustillo-Avendaño et al., 2018) and while tissue competence to generate these roots declines with age (Chen et al., 2014b; Xu et al., 2016), it can be restored by auxin application (Chen et al., 2014b).

A key remaining question is what factors control the timing and position of the auxin biosynthetic sources. Some regulators may be direct, such as the wound responsive transcription factor *ERF109* or the root maintenance and regeneration regulators *PLETHORA*, both shown to directly activate auxin biosynthesis (Kareem et al., 2015; Santuari et al., 2016; Bahieldin et al., 2018). Other regulators of auxin synthesis may be indirect, acting via the regulation of other hormones, such as cytokinin (Schaller et al., 2015), or by restricting the transcriptional response profile of the tissue. Indeed, as root tip regeneration process lasts 72h, during which multiple YUC enzymes are induced in a development-specific manner, it is unlikely that a single “regeneration factor” is responsible for activating auxin synthesis, but rather, a different network of redundant factors may act at different temporal windows.

### Auxin transport versus local biosynthesis in tip regeneration

Polar transport of auxin is a major regulator of auxin distribution within plant tissues (Adamowski and Friml, 2015). However, more recent evidence has suggested that local auxin biosynthesis may play at least an equal role in the process (Brumos et al., 2018; Zhao, 2018). Our work shows that during tip regeneration, local biosynthesis is the main regulator of auxin accumulation. This may be due to the small number of cells involved in regeneration or due to the need to form a pattern *de novo*. The latter hypothesis is more compelling, and indeed, recent work on the *de novo* formation of veins in leaves, thought to be mainly determined by PIN polarization (Scarpella et al., 2006), has shown that PINs may be dispensable for vein formation *per se* (Verna et al., 2019), while local auxin biosynthesis may play an important role (Kneuper et al., 2017; Ohashi-Ito et al., 2019). It should be noted that while PIN-mediated polar auxin transport may be redundant for regeneration, it does not follow that auxin transport is not a crucial player. Importantly, while auxin synthesis during regeneration occurs very close to the site of regeneration, the process was rescued by external hormone application, suggesting that biosynthesis may be required to establish local auxin concentrations on a tissue level, but some, non-PIN transport mechanisms may act to redistribute auxin inside relevant cells.

### A complex temporal sequence of hormone synthesis during regeneration

Plant development is governed by the relationship between different hormones and their relative activity in specific microenvironments (Schaller et al., 2015). Interestingly, application of a very low (5nM-10nM) auxin concentrations can restore regeneration capacity to L-Kyn treated, *ERF115:SRDX* plants or high cut plants. How can such a low hormone concentration induce this strong developmental effect? One possibility is that in the context of regeneration, auxin may promote its own synthesis, as observed during vasculature development (Ohashi-Ito et al., 2019), thereby generating a feedback loop that promotes the establishment of an auxin concentration peak near the cut site. It follows that factors controlling the sensitivity of cells to auxin may be just as important in allowing the injured tissue to generate sufficient auxin and promote successful regeneration. This view is supported by the reduced regeneration rates in mutants of the auxin response factor *MONOPTROS* (Efroni et al., 2016).

Plant regeneration and tissue reconstruction is a complex process which involves the activity of multiple factors and hormones. Our work shows that an injury-induced dynamic pattern of auxin sources, which likely integrates many developmental and environmental cues, is crucial for the decision of whether or not to regenerate the injured organ.

## Material and Methods

### Plant material and growth conditions

*yucQ* and ERF115-SRDX were previously described and characterized (Heyman et al., 2013; Chen et al., 2014a). The enhancer trap line J0571, *SCR:GFP, pDR5rev:3XVENUS-N7, TAA1p:GFP-TAA1, WOX5:mCherry WOL:GFP* and the SCR clone marker lines were previously described (Wysocka-Diller et al., 2000; Heisler et al., 2005; Stepanova et al., 2008; Efroni et al., 2016). *Arabidopsis* Col-O plants were grown on agar medium (0.5X Murashige and Skoog (MS), 0.5% sucrose, 0.8% agar), under long-day conditions (16h light and 8h dark), at 20°C, for 7 days. For NPA treatments, 7-day-old plants were transferred to agar plates supplemented with 10µM NPA. For L-Kyn (Sigma K8625) treatment, 7-day-old plants were transferred to agar plates supplemented with 100µM L-Kyn, unless indicated otherwise. For L-Kyn + IAA-treatmed plants were transferred to agar plates supplemented with 100µM L-Kyn and 100nM IAA. For split plate assays, plants were transferred to agar plates containing a barrier, with half of the plate containing 20µM L-Kyn and half containing 20µM L-Kyn and 10nM IAA.

### Root cutting assay and statistical analysis

7-day-old seedlings were cut 120µM or 220µM above the QC, according to (Sena et al., 2009). Between 2 and 4 independent cutting sessions (batches) were used for each experiment, each with a matching WT or mock control. To calculate statistical significance while accounting for batch effects, we used a logistic regression (models were Regenerated∼Genotype+Batch or Regenerated∼Treatment+Batch). Tukey HSD was used to derive the p-values for specific contrasts. To calculate batch-corrected regeneration rates, the regeneration rate of each treatment/genotype was normalized to the regeneration rate of WT/mock-treated plants of the same batch. All analyses were performed in R 3.5.2.

### Transgenic lines

All transgenic lines were generated using the golden gate cloning and the MoClo system (Engler et al., 2009) and were inserted to WT Col-O background. Promoter WOX5 was designed using the 4463bp fragment upstream to the first ATG of the *WOX5* gene. Promoter YUC9 was designed using the 3895bp fragment upstream to the first ATG of the *YUC9* gene. Promoter AHP6 was designed using the 5079bp fragment upstream to the first ATG of the *AHP6* gene. *amirYUC* was synthesized as a cistronic diMir, with two AtMir159a backbones. Sequence for *amirYUC* and primers are listed in Supplemental Table S2.

### Microscopy

Roots were observed using a Leica SP8 confocal microscope with x20 or x63 water objectives. Propidium iodide solution (0.01μg/ml) was used to stain the cell wall. A 552 nm and 488 nm lasers were used for excitation of GFP/YFP and mCherry/PI, respectively.

### Meristem isolation

Six-day old plants were either cut at their root tips or transferred whole to liquid protoplast solution (3% cellulose, 1% macerozyme, 0.4M mannitol, 20.48 mM MES, 0.02M KCl (1M), 0.1% BSA, 0.02 M CaCl2, pH 5.7) for a 10-15 minutes incubation period. The liquid solution caused the dissociation of the meristems at the transition zone. Meristems were then collected using a pipette, rinsed with DDW, and flash frozen, followed by RNA extraction using the QIAGEN RNeasy™ micro-kit according to the manufacturer’s protocol.

### RNA-seq analysis

Libraries were prepared using the 3’ mRNA-QuantSeq kit (Lexogen) according to the manufacturer’s instructions, followed by sequencing using NextSeq 500 (Illumina). Sequences were aligned to the TAIR10 genome using Bowtie2; gene expression hits were computed using htseq-count. To account for 3’ mis-annotation, the 3’ of all genes was extended 500bp downstream. Normalization and significance calling was performed using DeSeq2. Experiments were performed in 2, 3 or 5 replicates.

### qRT analysis

RNA was extracted using the RNeasy micro-kit (Qiagen), following by cDNA synthesis using qPCRBIO cDNA Synthesis Kit (PCR BIOSYSTEMS). Real-time PCR was performed using Fast SYBR® Green Master Mix (Rhenium) with a final primer concentration of 0.2μM. Primers are listed in Supplemental Table S2.

## Acknowledgments

We thank Jose Alonso, Anna Stepanova and Lieven de Veylder for sharing research material. We thank Yuval Eshed and Sigal Savaldi-Goldstein for comments and discussions. IE is supported by the Israeli Science Foundation (ISF966/17) and the Howard Hughes Medical Institute International Research Scholar Grant (55008730).

## Accession Numbers

RNA-seq data were deposited in GEO (GEO series).

## Supplemental Figures

**Supplementary Figure S1.**
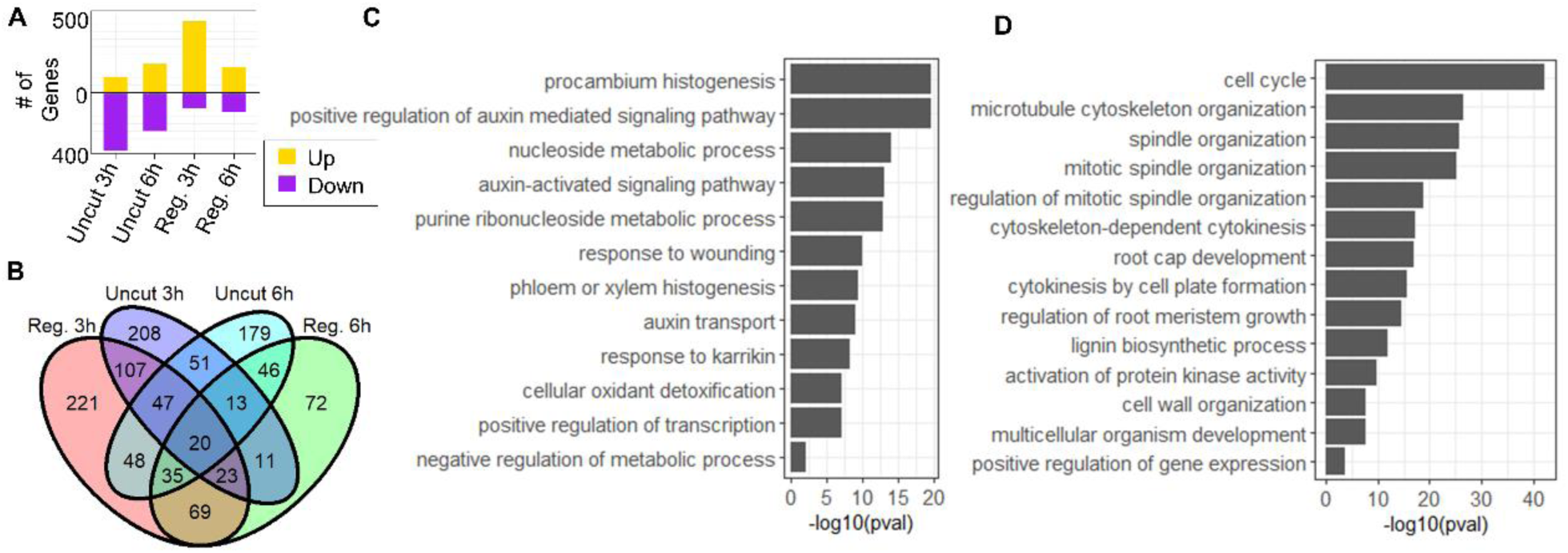
Analysis of gene expression changes in regenerating roots treated with L-Kyn. **A)** Number of genes whose expression was modified by L-Kyn in cut and uncut roots. **B)** Venn diagram of the modified genes of regenerating (Reg.) or uncut root meristems following 3h and 6h of L-Kyn treatment. Genes specifically modified by L-Kyn in regenerating roots, and used for downstream analysis. **C-D)** Enriched GO terms for genes suppressed by L-Kyn treatment in regenerating root tips at 3 hpc (C) and 6 hpc (D).

**Supplemental Figure S2.**
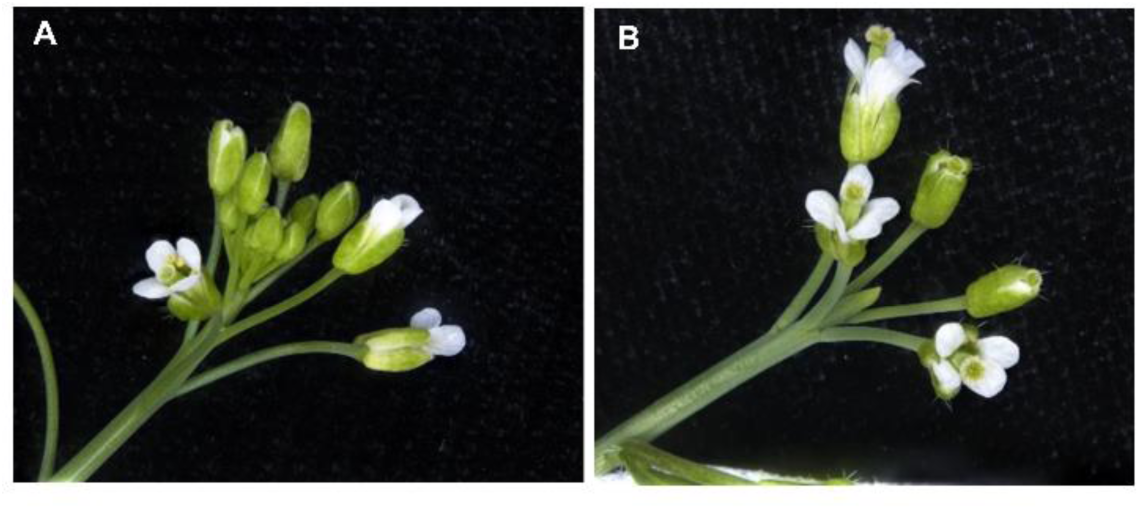
Strong lines of *pYUC9:amiRYUC* develop pin-like terminated meristems. **A)** WT floral meristem. **B)** *pYUC9:amiRYUC* pin-like floral meristem.

**Supplemental Figure S3.**
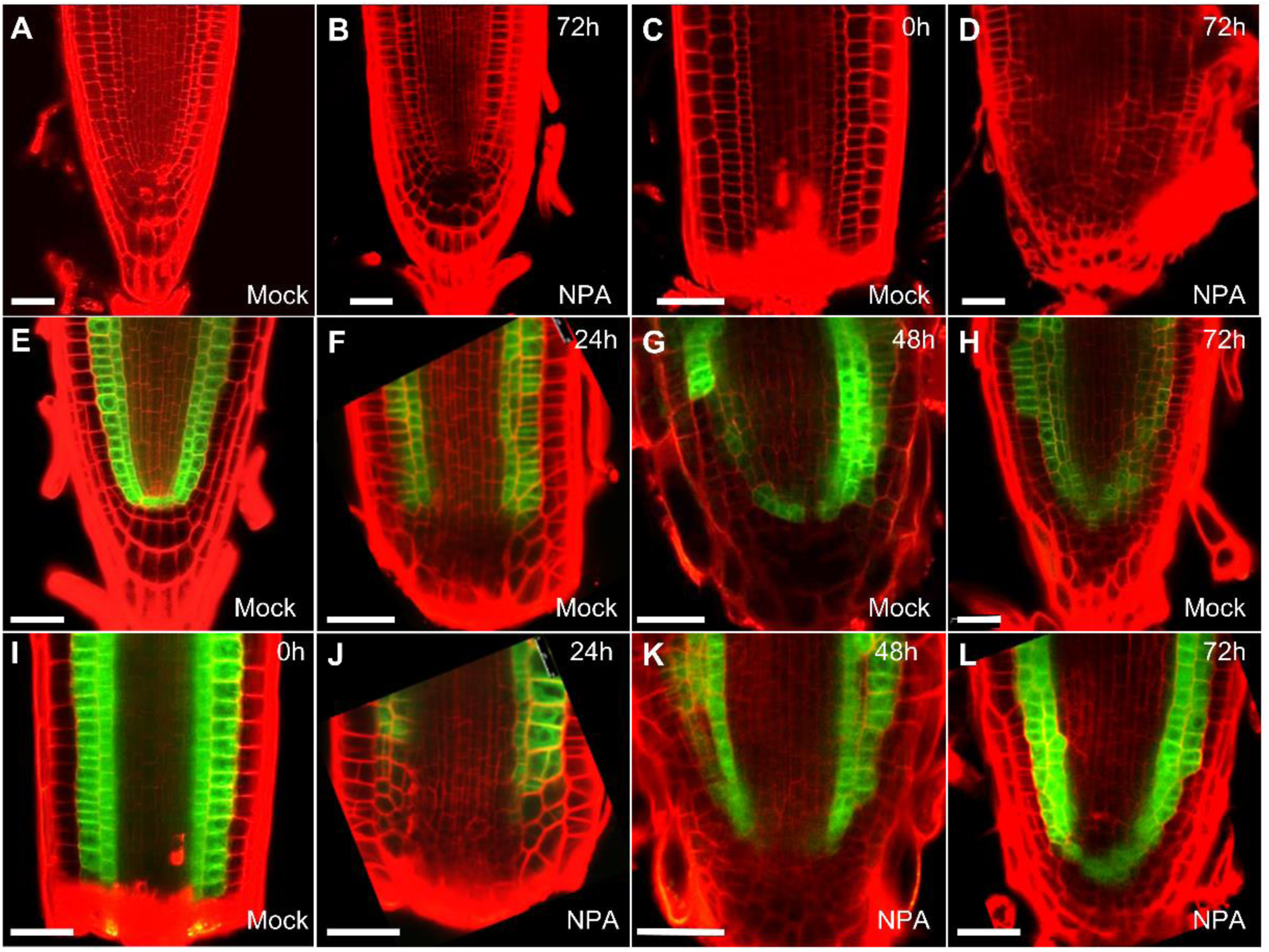
Regeneration and tissue pattern recovery in the presence of NPA. **A-D)** Confocal images of uncut (A-B) or cut (C-D) 7 DAS roots before (A,C) or after 72h of 10µM NPA treatment (B,D). **E-L)** Confocal images of the ground tissue marker J0571 in uncut (E) or regenerating (F-H, I-L) roots treated with mock (F-H) or 10µM NPA (J-L). Scale bar: 50µm.

**Supplementary Figure S4.**
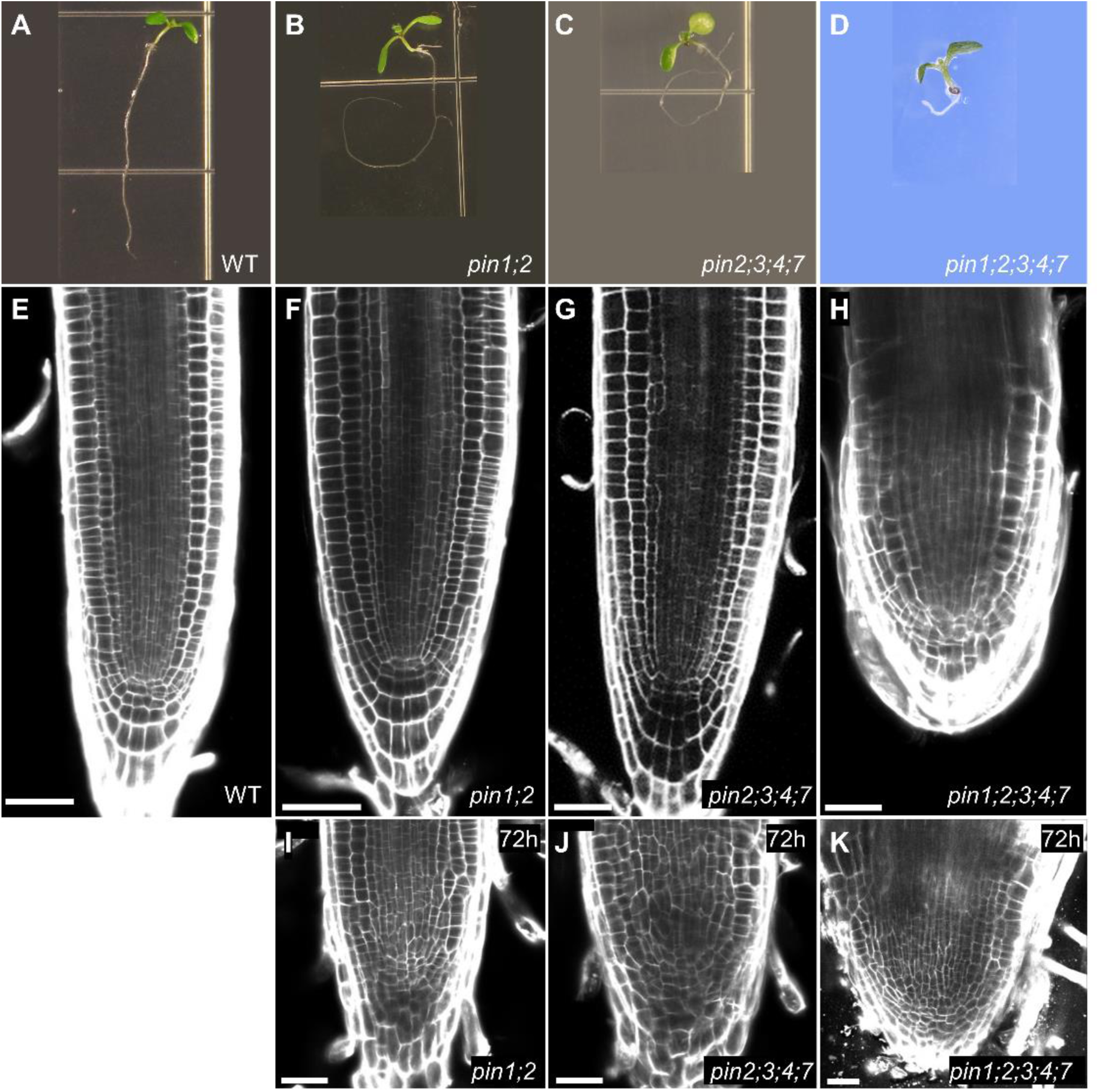
Root meristem growth, morphology and regeneration of high-order *pin* mutants. **A-D)** Images of 7 DAS WT (A) and *pin* mutants (B-D). **E-I)** Confocal images of uncut (E-H) or regenerating roots at 72 hpc (F-I) of WT (E) and *pin* mutants (F-K) Scale bars are 5mm in (A-D) and 50µm (E-K).

**Supplemental Figure S5.**
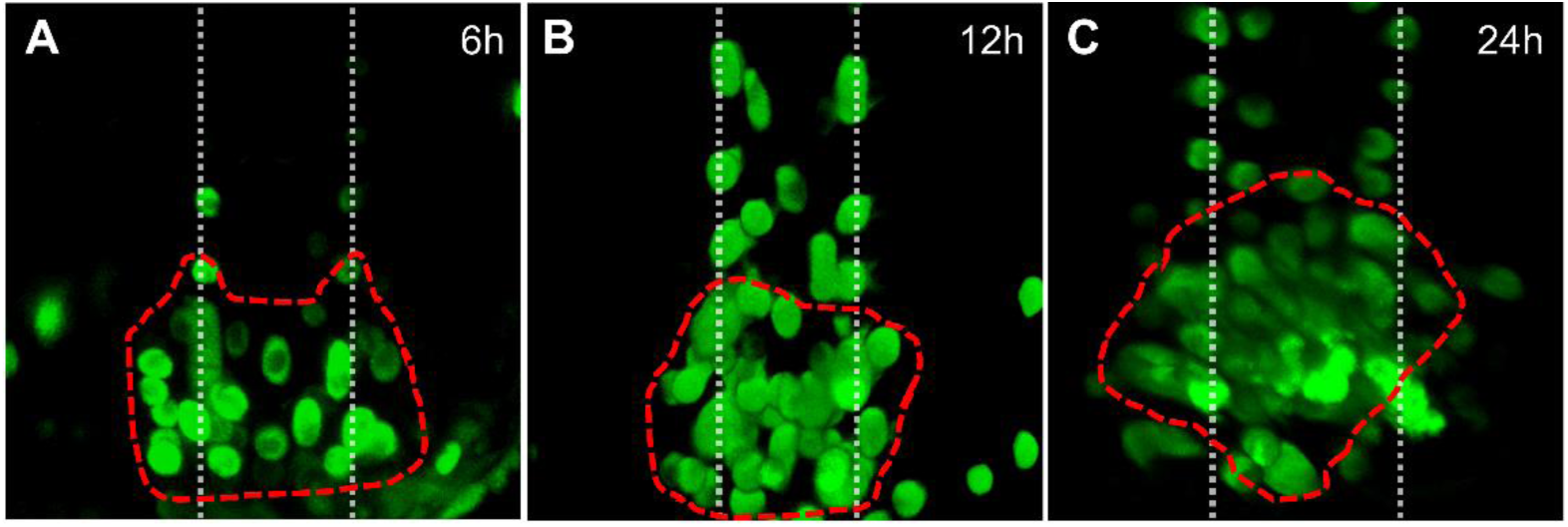
Regeneration of *big* mutants. **A-C)** Close-up of the regeneration region of *big DR5rev:3xVENUS-N7* plants during regeneration.

**Supplemental Table S1. Genes modified by L-Kyn at 3 h and 6h**.

**Supplemental Table S2. Primers and synthesized microRNAs used in this study**.

